# Souporcell3: Robust Demultiplexing for High-Donor Single-Cell RNA-seq Datasets

**DOI:** 10.1101/2025.07.10.664218

**Authors:** Minindu Weerakoon, Hai Vu, Reza Behboudi, Haynes Heaton

## Abstract

**Motivation:** Accurate demultiplexing of pooled single-cell RNA-seq (scRNAseq) data is critical for large-scale studies. However, existing methods like vireo, while effective up to ∼16 donors, often struggle with poor clustering due to local optima as donor numbers rise. In high-donor scenarios, overlapping genotypes, a dense genotype space, and increased doublet formation make demultiplexing challenging, requiring methods that are robust to sparse, high-dimensional data and maintain reliable accuracy even as sample complexity grows.

**Results:** We present an enhanced version of souporcell capable of demultiplexing up to 64 donors. The method uses 10x merge for initialization, K-Harmonic Means for robust clustering, and iterative refinement with reinitialization of low-quality clusters and locking of high-quality ones. Compared to vireo, vireo with overclustering, and the original souporcell, our approach completely eliminates duplicate clusters and achieves consistently high Adjusted Rand Index (ARI) scores across various doublet rates, demonstrating improved accuracy and scalability.

**Availability:** Souporcell3 source code and documentation are released on GitHub: https://github.com/wheaton5/souporcell

## 1. Introduction

Multiplexing cells from multiple individuals into a single scRNAseq sample has become a popular experimental design because it reduces costs, avoids batch effects, increases statistical power, improves doublet detection, and allows us to measure and remove ambient RNA(Heaton *et al*. 2020; Young and Behjati 2020). Demultiplexing these samples was originally enabled by demuxlet(Xu *et al*. 2019) which required a priori knowledge of the genotypes of every individual in the mixture. If this information was not available, additional costly DNA sequencing of each sample was required. Later, souporcell(Heaton *et al*. 2020) and vireo(Huang, McCarthy and Stegle 2019) were developed to demultiplex cells by genotype without prior knowledge of their genotypes using various sparse clustering methods. But clustering is known to be an NP-Hard problem to find the optimal clustering under any non-trivial loss function(Aloise *et al*. 2009). This is especially true as the number of clusters (or multiplexed samples) increases(Inaba, Katoh and Imai 1994; Mahajan, Nimbhorkar and Varadarajan 2012) with the historical solution to overcome local optima of doing multiple random restarts. It has been shown the number of clusters is in the exponent of the number of random restarts needed to even probabilistically find the global optima. More recently, the kmeans++(Arthur and Vassilvitskii 2006) cluster initialization strategy has provided a simple but strong heuristic method for finding initial values of cluster centers that are much more likely to converge to the global optimal in the cluster optimization process. Because kmeans++ uses individual data points as initial cluster centers, and new cluster centers are chosen randomly weighted on the squared distance to the closest existing cluster center, this poses two problems when dealing with sparse data. 1. It may be unclear what value to give the cluster center in dimensions for which the data point has no data. And 2. You can only judge the distance of a data point to the other cluster centers based on the dimensions for which it has data. 3. Because not every cell expresses every gene, the full gene is not covered by reads from each cell, and the experimental yield is not perfect, scRNAseq data has a sparsity of roughly 5% (only 5% of the dimensions have data for each cell). For this reason, kmeans++ is not suitable for scRNAseq data and other very sparse data types. Here we show multiple algorithmic improvements for clustering scRNAseq data by genotype. Previously, we showed souporcell had the ability to cluster up to ∼21 individuals with very high quality data but never recommended anyone design experiments with >16 individuals. Here, we show robust clustering by genotype up to 64 individuals. This will decrease costs while further increasing the statistical power of multiplexed scRNAseq experiments. Souporcell3 is freely available under the MIT open-source license at https://github.com/wheaton5/souporcell.

## 2. Methods

Souporcell3 introduces several key improvements over its predecessor souporcell(Heaton *et al*. 2020), enabling robust clustering of up to 64 donor samples. The clustering process begins with an 10x merge cluster center initialization strategy, in which the algorithm initializes ten times more clusters than the expected number of donors. Clusters that are in close proximity within the genotype space are iteratively merged until the desired number of clusters is reached. This approach helps capture subtle genetic variations and ensures that closely related clusters (those with a low number of differentiating alleles) are initially separated. (See supplementary 3 for a comparison of different cluster center initializing methods)

For the main clustering task, souporcell3 utilizes the K-Harmonic Means (KHM) algorithm. KHM is particularly effective in scenarios where poor cluster initialization can adversely affect outcomes, as it reduces sensitivity to initial conditions and enhances convergence to optimal solutions(Zhang, Hsu and Dayal 1999; Zhang 2001; Güngör and Ünler 2007, 2008). Further, souporcell3 uses a deterministic annealing variant similar to that of souporcell; however, souporcell3 applies it to KHM with a refined temperature constant. The temperature parameter starts high and gradually decreases, allowing the algorithm to explore globally optimal clusterings. (See supplementary 4 for a comparison of clustering methods with and without deterministic annealing).

After the initial clustering run, souporcell3 evaluates the quality of each cluster using both the number of cells assigned to the cluster and the associated loss value. Clusters that are identified as outliers, clusters deemed suboptimal, are reinitialized using the 10x merge cluster initialization strategy. Ten times the number of outlier clusters are initialized and then merged to generate well-separated cluster centers equal in number to the original outliers. This process allows the algorithm to improve clustering in the next iteration. At the same time, high-quality clusters are randomly selected from the remaining set, after excluding the outliers, and are locked to preserve their values during subsequent iterations. This strategy is inspired by node freezing and dropout in machine learning, which are used to reduce overfitting(Srivastava *et al*. 2014; Liu, Agarwal and Venkataraman 2021). This iterative refinement ensures accurate donor assignment and enhances the overall robustness of the clustering process (See supplementary 5 for more details).

## 3. Results

To assess the demultiplexing accuracy and robustness of our improved souporcell method, we constructed a benchmark dataset by integrating single-cell RNA-seq data from four different sources (Human Cell Atlas 2020a, Human Cell Atlas 2020b, Human Cell Atlas 2021, ENA 2020) (see Supplementary 1 for dataset composition and preprocessing details). This dataset was specifically designed to test performance under increasing levels of complexity, including doublet rates of 0%, 5%, and 10%. The results shown are using the 10% doublet rate 64-donor dataset, which is slightly higher than the expected 8% for a sample containing ∼20,000 cells. (10X Genomics 2020) (Fig. 1B, C).

**Figure 1.**
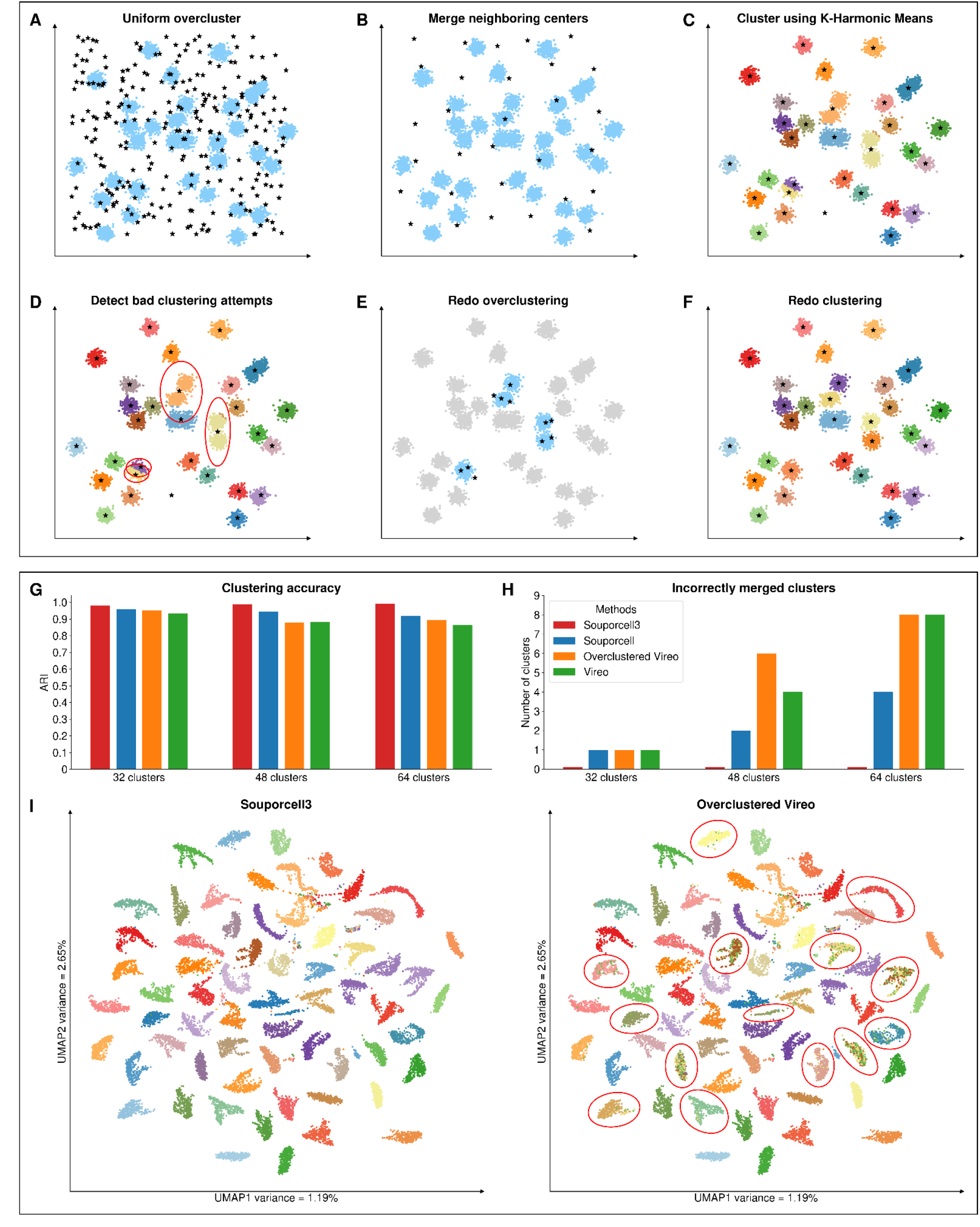
**A-F** Method overview: **A**. Randomly initialize with k x 10 clusters to ensure broad coverage of the genotype space. **B**. Merge nearby cluster centers using a distance metric defined as the sum of allele fraction differences, weighted by allele counts, until k clusters remain. **C**. Clustering using K-Harmonic Means (KHM) to ensure enhanced convergence and reduced sensitivity to initialization. **D**. Identify low and high quality clusters based on the number of assigned cells and the loss value per cluster. **E**. Reinitialize low-quality clusters **F**. Rerun KHM on the refined set while locking the high-quality clusters. **G-I** Results: **G**. ARI values for vireo, vireo with overclustering, souporcell, and souporcell3 for 10% doublet, 64-donor dataset. **H**. Number of incorrectly merged clusters for the same methods and dataset. **I**. Cluster maps using UMAPs for the x and y axes, showing the result of 10% doublet, 64-donor dataset after running souporcell3 (left) and vireo with overclustering (right) with poor clustering results highlighted with circles.

The UMAP plots in (Fig. 1I) display cells colored by their ground truth donor identity and project the high-dimensional clustering into two dimensions along the UMAP1 and UMAP2 axes, capturing the overall data structure. In the left plot, representing Souporcell3 clustering, clusters are well-separated across the space. Even in densely packed regions, such as the center, the clusters remain clearly partitioned, with only a few outliers, highlighting Souporcell3’s ability to resolve tightly grouped but distinct donor profiles. By contrast, the Vireo clustering shown in the right plot, exhibits several artifacts, including incorrectly merged clusters where cells from a single donor are split into multiple clusters (circled), indicating reduced clustering accuracy in complex settings.

We benchmarked our method against three state-of-the-art genotype-based demultiplexing approaches: vireo (using its default configuration), vireo with overclustering (where we specified the pre-cluster donors to k x 10 to match the number of clusters in our 10x merge), and the original souporcell (run with its default settings), while souporcell3 used the method described above. Clustering outcomes were evaluated using two primary metrics: the presence of incorrectly merged clusters after convergence (Fig. 1H), and clustering accuracy as measured by the ARI(Rand 1971; Hubert and Arabie 1985; Vinh, Epps and Bailey 2009) (Fig. 1G) (See supplementary 2 for explanation of the metrics and supplementary 6 for results on other datasets).

Incorrectly merged cluster analysis (Fig. 1H) shows that in high donor counts, our enhanced souporcell3 method produced no incorrectly merged clusters, in contrast to the comparison methods, which generated multiple such artifacts, particularly at higher donor counts. ARI analysis (Fig. 1G) further confirms the accuracy of our method, with consistently higher values across all conditions compared to the alternative methods. These results indicate the improvements introduced in our pipeline not only eliminate duplicates but also significantly enhance clustering fidelity.

Vireo with overclustering outperformed vireo with its default setting, highlighting that initializing cluster centers with additional pre-clustered donors yields better performance in high-donor scenarios. Souporcell, utilizing its expectation-maximization algorithm with random cluster center initialization, performed better than both versions of vireo which uses variational bayesian inference. Souporcell3 improves upon this by incorporating a multi-stage clustering pipeline that combines 10x merge initialization, K-harmonic means, and iterative cluster refinement with selective cluster locking. This accuracy gain remains robust even as data complexity and size increase, indicating that our method is well-suited for large-scale single-cell studies involving pooled donors.

## 4. Conclusion

Our evaluation of genotype-based demultiplexing using a complex dataset demonstrates that the improved souporcell3 approach consistently enhances clustering performance. The method scales effectively to 64 donors while maintaining high accuracy under varying levels of doublets and across different dataset complexities. By combining 10x merge initialization, K-Harmonic Means clustering, and iterative refinement with selective cluster locking, our method eliminates common artifacts such as empty or duplicate clusters and achieves higher concordance with ground truth donor labels. These enhancements highlight the advantages of robust initialization and adaptive cluster correction strategies in high-throughput demultiplexing. Future work will continue to refine the method in the context of different data types such as scATACseq and long read scRNAseq with even larger sample sizes and increasingly complex single-cell datasets, ensuring accurate and scalable demultiplexing for population-scale single-cell studies.

## Acknowledgement

None

## Supplementary Data

Link **Funding** None

## Data availability

Souporcell3 source code and documentation are released on GitHub: https://github.com/wheaton5/souporcell

The single-cell RNA-seq datasets used and analyzed in this study are publicly available from the Human Cell Atlas under the following project accessions: 3089d311-f9ed-44dd-bb10-397059bad4dc, cc95ff89-2e68-4a08-a234-480eca21ce79, and fcaa53cd-ba57-4bfe-af9c-eaa958f95c1a.

European Nucleotide Archive (ENA) data used are available under sample accessions ERS2630502–ERS2630507, corresponding to the cell lines euts, nufh, babz, oaqd, and ieki.

