## supplementary for "Souporcell3: Robust Demultiplexing for High-Donor Single-Cell RNA-seq Datasets"

### 1. Dataset

#### 1.1 Data used

We utilized four datasets from the Human Cell Atlas and European Nucleotide Archive, each comprising human-derived samples and generated using the 10X Genomics 3' chemistry (v1 and v2). All samples were derived from Homo sapiens, and the data collectively span a wide range of immune and developmental contexts, offering a diverse foundation for evaluating demultiplexing performance across donor pools and biological conditions.

Source 1

Publication: Immune Landscape of Viral- and Carcinogen-Driven Head and Neck Cancer (Cillo et al. 2020)

Donors: 37

Cell count: 126.0k

Dataset: (Human Cell Atlas 2020a)

Source 2

Donors: 28

Cell count: 1.5M

Dataset: (Human Cell Atlas 2020b)

Source 3

Publication: Mapping the developing human immune system across organs (Jardine et al. 2021)

Donors: 30

Cell count: 922.3k

Dataset: (Human Cell Atlas 2021)

Source 4

Donors: 5

Cell count: 50k

Dataset: (ENA 2020)

Collectively, these datasets span 100 unique donors and over 2.5 million single-cell transcriptomes, providing a rich and diverse foundation for evaluating and benchmarking single-cell demultiplexing methods in real-world, heterogeneous biological contexts.

#### 1.2 Donor allocation

Out of the initial 100 unique donors, we filtered out samples with corrupted data, insufficient read depth, and redundant samples from the same individual collected across different organs. After applying these quality control and deduplication steps, we curated a high-quality dataset comprising 64 distinct donors. The table below summarizes the composition of these 64 donors and their respective sources.

**Table 1.** Shows the source and the original ID of the modified ID used in the CB tag of our dataset for differentiating donors.

| Modified ID | Original ID | Source | Modified ID | Original ID | Source |
|---|---|---|---|---|---|
| 0 | euts | ENA 2020 | 35 | CB6 | Human Cell Atlas 2020b |
| 1 | BL1 | Human Cell Atlas 2020b | 36 | ieki | ENA 2020 |
| 2 | CB2 | Human Cell Atlas 2020b | 37 | F38 | Human Cell Atlas 2021 |
| 3 | HNSCC_9 | Human Cell Atlas 2020a | 38 | F45 | Human Cell Atlas 2021 |
| 4 | HNSCC_3 | Human Cell Atlas 2020a | 39 | F32 | Human Cell Atlas 2021 |
| 6 | HD_5 | Human Cell Atlas 2020a | 41 | BL7 | Human Cell Atlas 2020b |
| 7 | HNSCC_20 | Human Cell Atlas 2020a | 42 | CB4 | Human Cell Atlas 2020b |
| 8 | HD_tonsil2 | Human Cell Atlas 2020a | 43 | F30 | Human Cell Atlas 2021 |
| 9 | BM8 | Human Cell Atlas 2020b | 44 | HNSCC_14 | Human Cell Atlas 2020a |
| 10 | HNSCC_18 | Human Cell Atlas 2020a | 45 | BM4 | Human Cell Atlas 2020b |
| 11 | BM2 | Human Cell Atlas 2020b | 46 | CB11 | Human Cell Atlas 2020b |
| 12 | BM5 | Human Cell Atlas 2020b | 47 | F23 | Human Cell Atlas 2021 |
| 14 | CB10 | Human Cell Atlas 2020b | 48 | F29 | Human Cell Atlas 2021 |
| 15 | HD_2 | Human Cell Atlas 2020a | 49 | HD_3 | Human Cell Atlas 2020a |
| 16 | CB5 | Human Cell Atlas 2020b | 50 | HNSCC_21 | Human Cell Atlas 2020a |
| 17 | BL6 | Human Cell Atlas 2020b | 51 | HD_4 | Human Cell Atlas 2021 |
| 18 | nufh | ENA 2020 | 52 | BM3 | Human Cell Atlas 2020b |
| 19 | BL8 | Human Cell Atlas 2020b | 53 | HNSCC_13 | Human Cell Atlas 2020a |
| 20 | CB1 | Human Cell Atlas 2020b | 54 | HNSCC_19 | Human Cell Atlas 2020a |
| 21 | BL2 | Human Cell Atlas 2020b | 55 | F61 | Human Cell Atlas 2021 |
| 22 | F51 | Human Cell Atlas 2021 | 56 | CB9 | Human Cell Atlas 2020b |
| 23 | HNSCC_23 | Human Cell Atlas 2020a | 57 | HNSCC_2 | Human Cell Atlas 2020a |
| 25 | BM1 | Human Cell Atlas 2020b | 58 | CB3 | Human Cell Atlas 2020b |
| 26 | HNSCC_11 | Human Cell Atlas 2020a | 59 | CB7 | Human Cell Atlas 2020b |
| 27 | HNSCC_16 | Human Cell Atlas 2020a | 60 | BL4 | Human Cell Atlas 2020b |
| 28 | BM6 | Human Cell Atlas 2020b | 61 | HNSCC_6 | Human Cell Atlas 2020a |

| | | | | | |
|---|---|---|---|---|---|
| 29 | F19 | Human Cell Atlas 2021 | 62 | oaqd | ENA 2020 |
| 30 | HNSCC_24 | Human Cell Atlas 2020a | 63 | BM7 | Human Cell Atlas 2020b |
| 31 | HD_1 | Human Cell Atlas 2020a | 64 | CB12 | Human Cell Atlas 2020b |
| 32 | F21 | Human Cell Atlas 2021 | 66 | HNSCC_22 | Human Cell Atlas 2020a |
| 33 | HNSCC_7 | Human Cell Atlas 2020a | 67 | babz | ENA 2020 |
| 34 | BL5 | Human Cell Atlas 2020b | 69 | BL3 | Human Cell Atlas 2020b |

#### 1.3 Preprocessing

From the collected datasets, we generated subsets tailored for benchmarking by extracting the required number of cells for each experimental scenario. Specifically, we constructed three datasets with controlled doublet rates of approximately 0%, 5%, and 10%. Each subset was limited to a total of around 10,000~30,000 cells, aligning with the practical upper limit for many current single-cell RNA-sequencing (scRNA-seq) experiments using 10X Genomics platforms (10X Genomics 2020). This cell count threshold reflects a balance between throughput and data quality, as higher cell input often increases the likelihood of doublet formation and misclassification errors. Published reports suggest that, at this scale, the expected doublet rate for standard 10X Chromium 3' chemistry ranges between 5% and 10% when targeting ~20,000 cells (10X Genomics 2020; Zheng et al. 2017; Wolock et al. 2019; McGinnis et al. 2019).

**Table 2.** Shows the composition of different datasets used in evaluation.

| Type | Donors | Total cell count | Cells per donor | Doublet count | Doublet percentage |
|---|---|---|---|---|---|
| No doublets | 4 | 9300 | 2325 | 0 | 0 |
| | 8 | 10000 | 1250 | 0 | 0 |
| | 16 | 12800 | 800 | 0 | 0 |
| | 32 | 12800 | 400 | 0 | 0 |
| | 48 | 19200 | 400 | 0 | 0 |
| | 64 | 25600 | 400 | 0 | 0 |
| 5 percent doublet | 4 | 7123 | 1781 | 373 | 5.23655763 |
| | 8 | 8303 | 1038 | 428 | 5.154763339 |
| | 16 | 11200 | 700 | 578 | 5.160714286 |
| | 32 | 11777 | 368 | 617 | 5.239025219 |
| | 48 | 18239 | 380 | 959 | 5.257963704 |
| | 64 | 20061 | 313 | 1053 | 5.248990579 |
| 10 percent doublet | 4 | 6740 | 1685 | 740 | 10.97922849 |

| | | | | | |
|---|---|---|---|---|---|
| | 8 | 8120 | 1015 | 888 | 10.93596059 |
| | 16 | 10799 | 675 | 1199 | 11.1028799 |
| | 32 | 11159 | 349 | 1239 | 11.10314544 |
| | 48 | 16917 | 352 | 1877 | 11.09534787 |
| | 64 | 22679 | 354 | 2519 | 11.10719168 |

#### 2. Evaluation metrics: Adjusted Rand Index (ARI) and Incorrectly Merged Clusters (IMC)

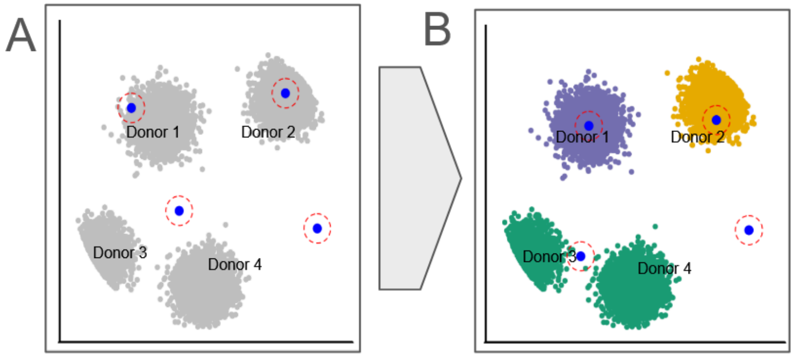


**Figure 1. A.** Before clustering, cluster centers are initialized **B.** After the clustering method converges, an empty cluster center is present (not assigned to any cells) and one cluster center is assigned to cells from two donors.

Throughout the main text and supplementary materials, we evaluate clustering performance using two key metrics: the Adjusted Rand Index, or ARI, and the number of Incorrectly Merged Clusters, or IMC. The Rand Index is a standard and widely used metric in clustering analysis; it measures the similarity between two data clusterings (Rand 1971). A more advanced method, known as the Adjusted Rand Index, is introduced to account for the random chance the clustering gets assigned correctly (Hubert et al. 1985). In our case, ARI is used, and we compare the clustering output of each method against the known ground truth, which is available since the datasets were generated from sources with known donor origins. The second metric, IMC, captures instances where two or more distinct ground-truth donors are incorrectly assigned to the same cluster by the method. In figure 1B, the IMC value is one, as both donor 3 and donor 4 cells are incorrectly grouped into the same cluster by the clustering algorithm.

#### 3. Cluster Initialization

##### 3.1 *k*-means++ initialization

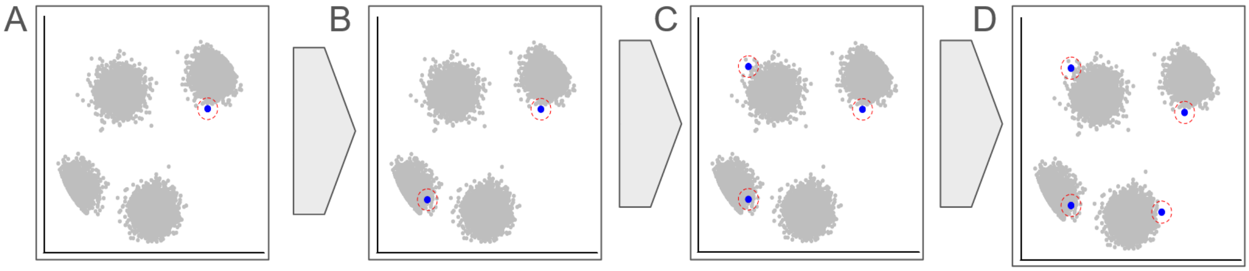


**Figure 2. A.** Randomly select a cell to be the first cluster center **B, C, D.** Iteratively select cells as cluster centers based on their squared distance to the closest existing cluster center.

Almost all commonly used clustering algorithms implicitly assume the data is dense and continuous, which ensures a relatively smooth cost function landscape. In contrast, genetic data is both sparse and discrete, with each dimension typically limited to three possible values. This sparsity poses significant challenges for greedy clustering algorithms like *k*-means, which are prone to getting trapped in local minima. A standard remedy is to perform multiple random restarts; however, the number of restarts needed to reliably approach the global optimum increases exponentially with the number of clusters *k* (Aloise et al. 2009, Inaba et al. 1994). To address this, the *k*-means++ initialization method was proposed (Arthur & Vassilvitskii 2007), offering a simple, parameter-free, and generally robust strategy. Despite its widespread adoption, *k*-means++ still struggles in sparse settings, where limited shared information between data points across dimensions undermines the effectiveness of its distance-based seeding.

In *k*-means++, the initial cluster center is selected randomly from the dataset. A distance metric is then used iteratively to determine the probability of selecting a new cluster center, based on the weighted squared distance of each datapoint from its closest existing cluster center. For this metric, we used the beta-binomial distance, as it accounts for genotype positions where no data is present.

#### 3.2 Overclustering initialization

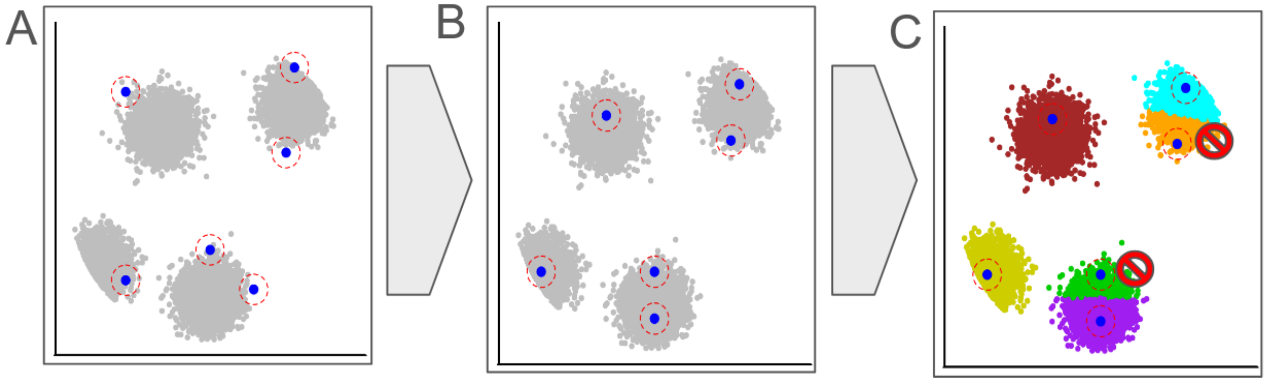


**Figure 3. A.** Selecting $k + \sqrt{k}$ random cells as cluster centers, where $k$ is the desired number of clusters **B.** Clustering using the selected cluster centers **C.** Select $k$ cluster centers with maximally associated cells.

Another strategy aimed at improving cluster initialization is overclustering. It is suited for challenging genetic datasets, such as those that are sparse and discrete. Traditional clustering algorithms often struggle in these contexts due to poor initial seeding, which can lead to convergence on suboptimal solutions. By temporarily increasing the number of clusters beyond the desired value, overclustering aims to better capture the underlying structure of the data, especially since the effective number of clusters can be higher than expected due to the presence of doublets. This strategy is employed by Vireo and SC3 (Huang et al. 2019, Kiselev et al. 2017) to enhance cluster initialization.

We begin clustering with $k + \sqrt{k}$ clusters instead of the target $k$. After performing the clustering step, we evaluate the resulting clusters based on the number of cells assigned to each. The $k$ clusters with the highest cell counts are then selected to form the final set of cluster centers, similar to Vireo. This selection criterion ensures that the most representative and well-supported clusters are retained, while smaller, potentially spurious clusters are discarded. However, this method can be prone to errors in high-donor-count scenarios, where the genotype space is limited and substantial overlap between donors exists. In such cases, selecting clusters solely based on cell count may result in duplicate donors, where one cluster gets assigned cells of multiple donors.

##### 3.3 10× merging initialization

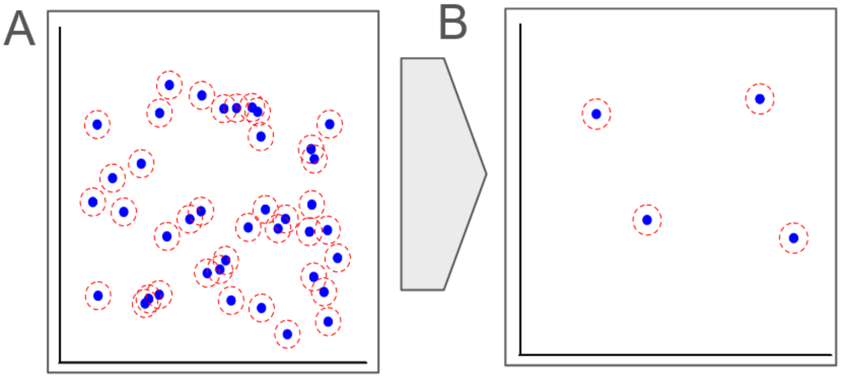


**Figure 4. A.** Randomly initializing 10× the number of *k* clusters required **B.** Merging clusters in close proximity with each other until required cluster size reached.

10× merging is the third and primary initialization strategy employed by Souporcell3, where the algorithm generates a large number of preliminary cluster centers—ten times the expected number of donors. Rather than using these centers directly for final clustering, the algorithm performs a merging step at first, combining those that are highly similar in genotype space—typically differing by only a few informative alleles—into a more concise and representative set. Using a distance metric—the sum of squared differences between randomly assigned cluster center values, weighted by the number of alleles at each variant position—we identify the two closest cluster centers and merge them (by calculating the mean values for each variant position and using that as the new cluster center). This process is repeated iteratively until the desired *k* cluster centers are reached.

10× merging reduces redundancy while preserving essential genotype distinctions, ensuring that the actual clustering step begins with informative and well-spread cluster centers. This is particularly helpful in scenarios where multiple overlapping donor cells occupy a dense genotype space, as widely separated cluster centers are unlikely to claim cells from multiple donors at the outset.

##### 3.4 Cluster initialization evaluation

To evaluate the different cluster initialization methods, we tested them on our 64-donor dataset without doublets. We also included an optimal cluster initialization blueprint, where clusters were initialized using ground truth donor information, in order to assess whether perfect initialization alone could give perfect clustering or if additional refinement steps were still necessary in high-donor scenarios.

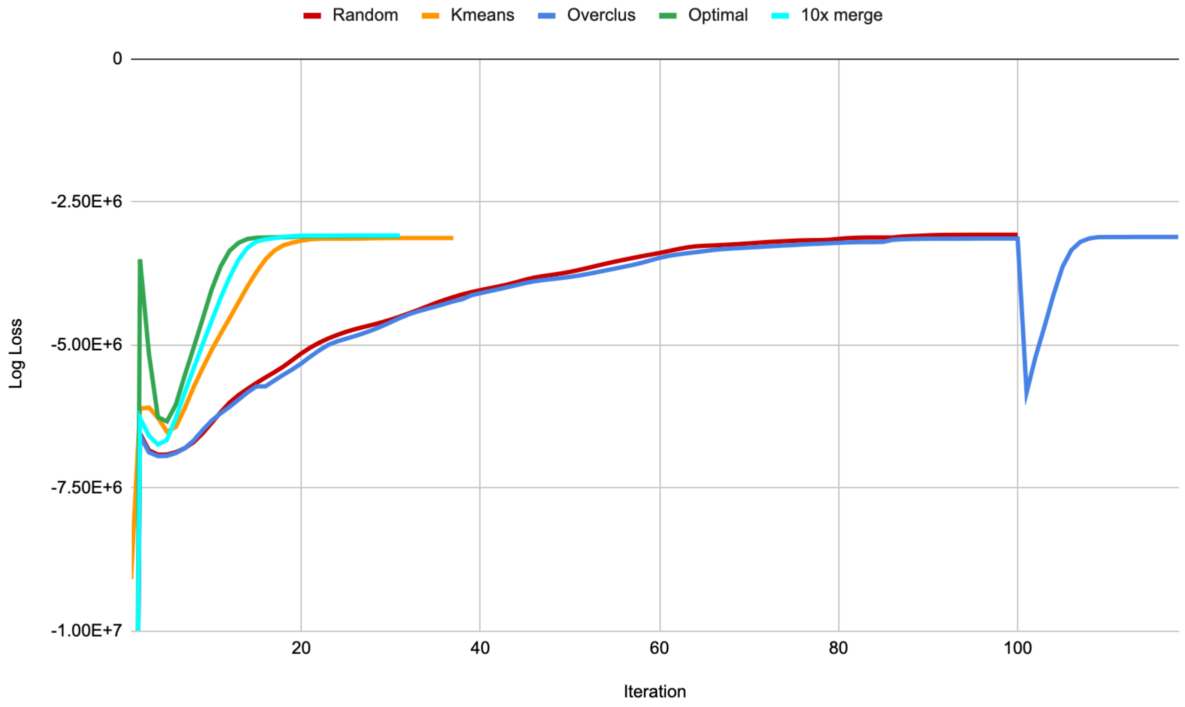


**Figure 5.** Logistic loss vs. iterations for EM with 5 different initialization methods: Random, Kmeans, Overclus, Optimal, and 10× merge. A no-doublet dataset with 64 donors was used for the experiment.

The logistic loss – number of iterations relationship for a no-doublet dataset of 64 donors is shown in the figure. Expectation-maximization (EM) with Optimal cluster initialization started off with the lowest log loss but eventually reached the same log loss as Kmeans and 10× merge, showing the same pattern. Both Random and Overclus methods took more iterations to converge, but Overclus ran for more iterations after removing the extra $\sqrt{k}$ clusters (100+ iterations) and converged to the same log loss as Random.

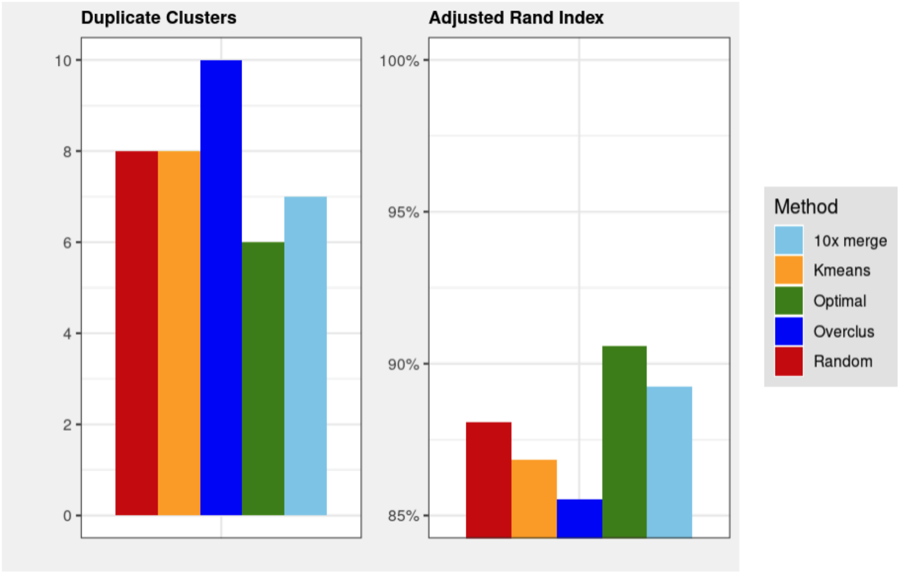


**Figure 6.** The right side shows the ARI values for an EM run with different cluster initialization methods. The left side illustrates the duplicate assignments present for the methods. The dataset used for figure 5 was also utilized for this evaluation.

The bar charts display two key clustering metrics: duplicate clusters (or IMC) and ARI, across five initialization methods: 10× merge, Kmeans, Optimal, Overclus, and Random. As shown on the left, the number of duplicate clusters is lowest for the optimal initialization method, which is expected since the cluster centers were initialized with ground truth values. Overclus and random initializations resulted in the highest number of duplicate clusters, with overclus making 10. Among the practical initialization methods, 10x merge performed the best, producing around 7 duplicates. On the right, the ARI values reinforce this trend, with the optimal method achieving the highest accuracy (> 90%), while overclus and random show lower performance (< 88%).

Interestingly, when using the optimal initialization, which uses ground truth information to set the cluster centers, the Rand index improved only marginally by 1.3%, and duplicate assignments were reduced by just one, when compared to 10x merge. This is unexpected, as perfect clustering is theoretically possible under such conditions, where no further updates to the cluster centers should be needed. However, even after convergence, duplicate assignments still occurred. This may be due to the large number of donors involved, where cells are densely packed in the genotype space. As the optimization progresses, log loss updates may inadvertently shift a cluster center toward a dense region, causing it to claim ownership of cells from neighboring donors with similar genotypes, thus introducing duplicates.

#### 4. Clustering

We showed that the high-donor clustering problem cannot be solved alone with cluster initialization; we try exploring the KHM clustering method to determine whether any improvements can be made.

##### 4.1 K-Harmonic Means clustering

Soft clustering methods, such as Expectation-Maximization (EM) applied to Gaussian Mixture Models (GMMs), have been widely used due to their ability to recover more optimal clusterings than hard clustering methods like Lloyd's algorithm for k-means (Peizhuang 1983). However, these methods are more prone to converging to local optima if the cluster initialization is not done correctly. To address this, we adopt K-Harmonic Means (KHM) (Zhang, Hsu & Dayal 1999, Güngör & Ünler 2007, Güngör & Ünler 2008), a soft clustering objective that computes the contribution of each data point to all cluster centers based on inverse distances. This formulation allows every point to influence all centers during each iteration, leading to more robust convergence and improved clustering outcomes. KHM has been shown to outperform traditional k-means in both optimality and resilience to poor or adversarial initializations. To further adapt KHM to high-dimensional genotype space, we employ a generalized variant, KHMp (Zhang 2001), which uses Lp-norm normalization to address symmetry issues known to affect clustering performance in such contexts. This introduces a tunable hyperparameter, p, that controls the influence of distance magnitudes. We found that setting p = 25 minimized the loss across a range of tested values, and this was used in all reported results.

##### 4.2 Deterministic annealing

Souporcell used a deterministic annealing variant of expectation–maximization (DA_EM) to manage non-convexity via a "temperature" parameter that starts high and is gradually lowered; this approach allows the solution to more often fall into the global optimal clustering. Similarly, souporcell3 employs a deterministic annealing-enhanced KHM (DA_KHM), combining the robustness of KHM with the annealing strategy: The temperature parameter is slightly adjusted using a temperature constant to account for the difference in convergence speed between the two clustering algorithms. When the temperature reaches 1.0, the loss function effectively becomes the KHM loss function. We compare the deterministic annealed methods and the vanilla methods below.

##### 4.3 K-Harmonic Means evaluation

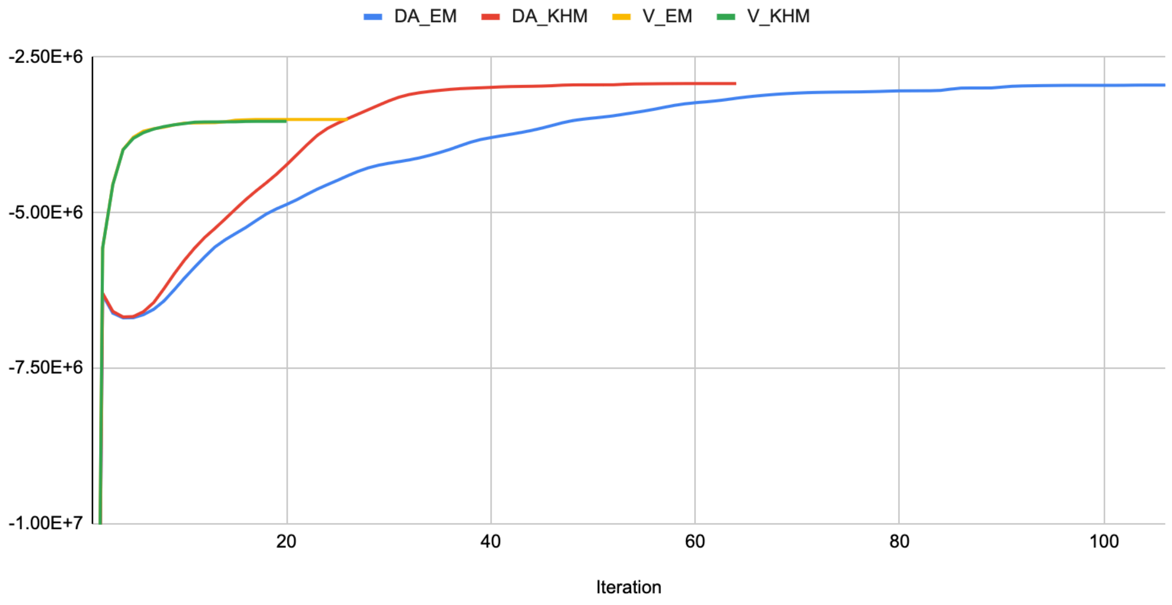


**Figure 7.** Log loss vs. iterations for clustering methods: DA_EM (Expectation maximization with deterministic annealing), DA_KHM (K Harmonic means with deterministic annealing), V_EM (Expectation maximization without deterministic annealing), and V_KHM (K Harmonic means without deterministic annealing). A no-doublet dataset with 64 donors was used.

The log loss versus iteration for a no-doublet dataset of 64 donors is shown in the figure. EM and KHM without deterministic annealing (V_EM and V_KHM) followed a similar pattern and converged to approximately the same log loss. Both deterministic annealing methods required more iterations to converge, with DA_EM taking about twice as many iterations as DA_KHM. DA_EM converged to a slightly worse log loss of -2,951,437.3 compared to -2,927,414.5 for DA_KHM.

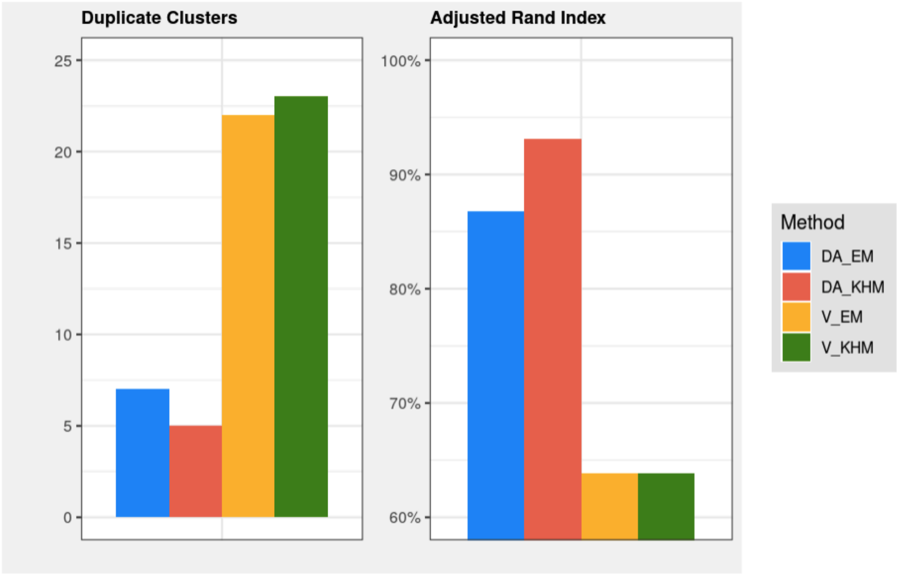


**Figure 8.** The right side shows the Adjusted Rand Index values for clustering methods DA_EM (Expectation maximization with deterministic annealing), DA_KHM (K Harmonic means with deterministic annealing), V_EM (Expectation maximization without deterministic annealing), and V_KHM (K Harmonic means without deterministic annealing). The left side illustrates the duplicate assignments present in the same methods. A no-doublet dataset with 64 donors was used for the evaluation.

The bar plots present the performance of four clustering methods, DA_EM, DA_KHM, V_EM, and V_KHM, on a no-doublet dataset of 64 donors. The left panel shows the number of duplicate clusters, and the right panel displays the Adjusted Rand Index (ARI), which reflects clustering accuracy. Among the methods, DA_KHM achieved the highest ARI, exceeding 93 percent, and also had the fewest duplicate clusters, just five. DA_EM also performed well, with an ARI close to 90 percent and only seven duplicate clusters. In contrast, both non-annealing variants (V_EM and V_KHM) performed significantly worse, with ARI values around 63 percent and more than 22 duplicate clusters each. These results demonstrate the substantial benefit of deterministic annealing in clustering complex genotype data.

The more robust DA_KHM method outperformed DA_EM by 5% in ARI and produced 2 fewer duplicate clusters. While this represents a significant improvement, perfect clustering is still not achieved, as some duplicate assignments remain.

#### 5. Rerun with cluster reinitialization

##### 5.1 Bad cluster center detection

We analyzed the cell-to-cluster assignments in the 64-donor, no-doublet dataset after algorithm convergence to investigate our hypothesis regarding duplicate assignments, specifically that cluster centers may incorrectly claim cells from other donors due to the constrained genotype space, and to develop a potential solution.

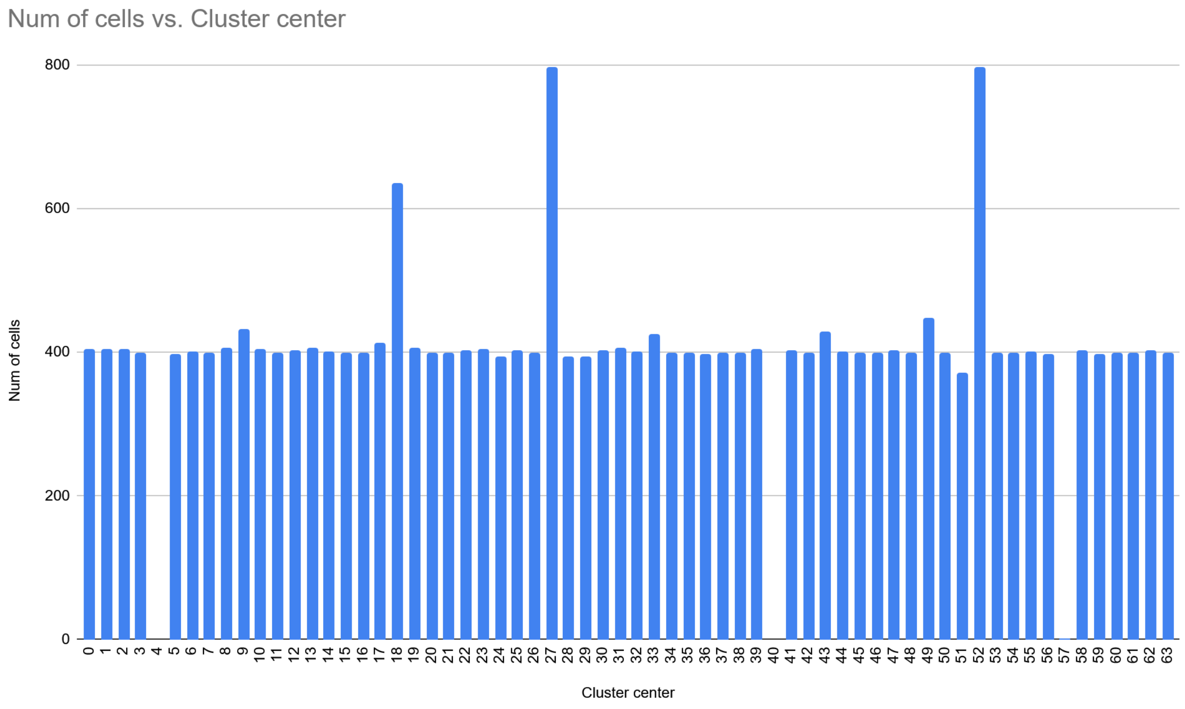


**Figure 9.** Number of cells assigned to each cluster center. A no-doublet dataset with 64 donors was used.

Analyzing the clustering output, we found that the number of duplicate clusters corresponds to empty or near-empty clusters. As shown in the figure, the cell assignments to each cluster center after convergence reveal three duplicate cluster centers, which align with the presence of three empty clusters.

By computing the total log loss for each cell's assigned cluster, normalizing it by the mean total log loss, and identifying statistical outliers, we can effectively pinpoint cluster centers that are poorly or incorrectly assigned.

$$CC\,Value = \sum_{all\ cells} \frac{Log\ loss\ for\ assigned\ CC}{Mean\ log\ loss}$$

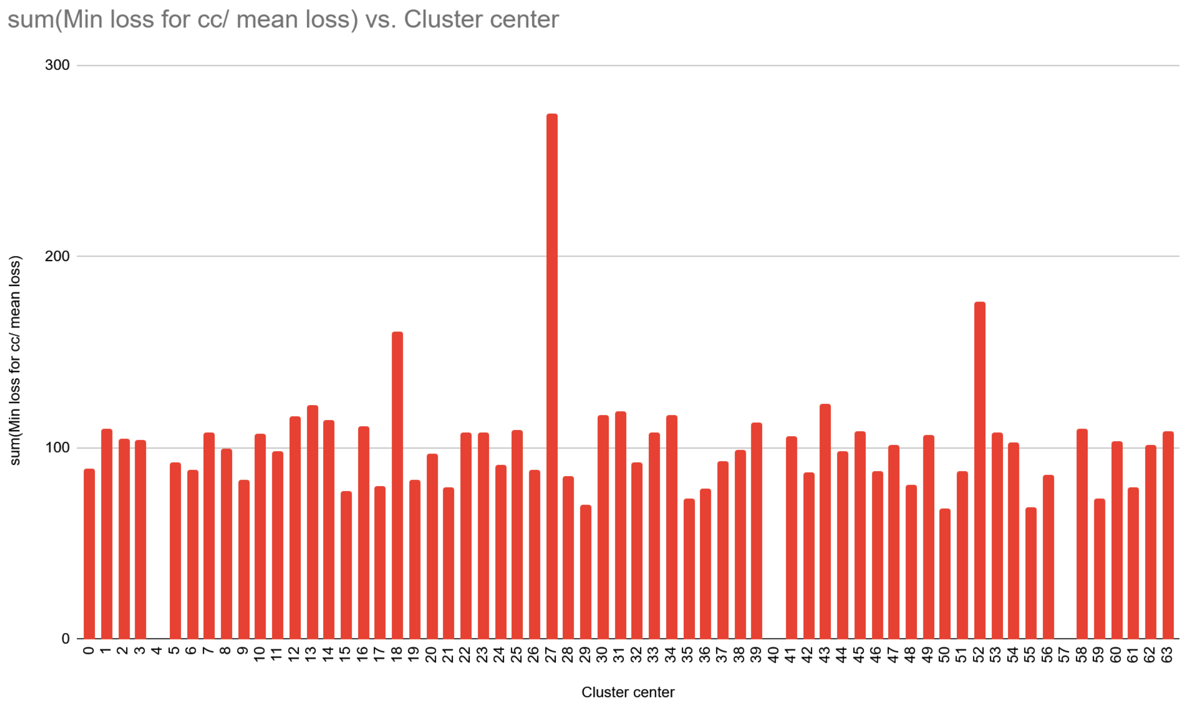


**Figure 10.** Total log loss for each cell's assigned cluster, normalized it by the mean total log loss. A no-doublet dataset with 64 donors was used.

The outliers with a high number of cell assignments (greater than the median plus 1.5 times the interquartile range) can be considered the culprits that claimed ownership of the low-assignment outliers (less than the median minus 1.5 times the interquartile range). By reinitializing these cluster centers, we should be able to achieve improved clustering results.

In scenarios where the distribution of cells per donor is uneven, it might not be possible to detect outliers using this method. However, empty or near-empty clusters can still be identified by defining an appropriate threshold for cell assignment counts. The corresponding duplicate clusters typically have cc values on the high end. In such cases, we select a number of high-loss clusters equal to the number of empty clusters and treat them as outliers for reinitialization. Even if a correctly clustered center is mistakenly selected as an outlier, the subsequent clustering should not significantly affect the final outcome, as it should be reassigned the same cells.

#### 5.2 Multiple runs with reinitialization and locking.

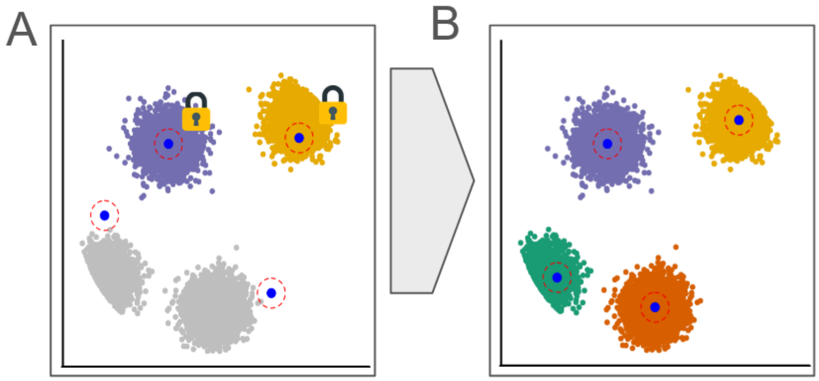


**Figure 11. A.** After the first clustering run, the bad clusters are detected and reinitialized while good clusters are locked **B.** While the good clusters are locked to updates the KHM clustering is converged.

Now that we can identify poorly formed clusters after algorithm convergence, we reinitialize the corresponding cluster centers and rerun the algorithm to improve clustering accuracy. Since most donors are correctly clustered in the initial run, it is unnecessary to update all the cluster centers. Instead, we lock the cluster centers deemed high quality, effectively freezing them, while allowing only the reinitialized (bad) clusters to update. This strategy is analogous to node freezing and node dropout in machine learning, where parts of a pretrained model remain fixed while selected components are fine-tuned on new data. (Srivastava et al. 2014, Ma et al. 2021). This process can be iteratively repeated, detecting bad clusters and reapplying the locked KHM algorithm with reinitialized centers, until no bad clusters are identified.

#### 6 Results

In this section, we present the results of different methods: vireo, vireo overclus, souporcell, and souporcell3. Vireo is run using its default configuration. For vireo overclus, we specify the pre-clustered donors to $10 \times k$ to match the cluster number in our 10x merge setting. Souporcell is run with its default settings. Souporcell3 uses the 10x merge cluster initialization method, the KHM clustering method, and is rerun with reinitialized bad and locked cluster centers. The datasets used are the ones described in supplementary 1.

#### 6.1 Duplicate cluster centers

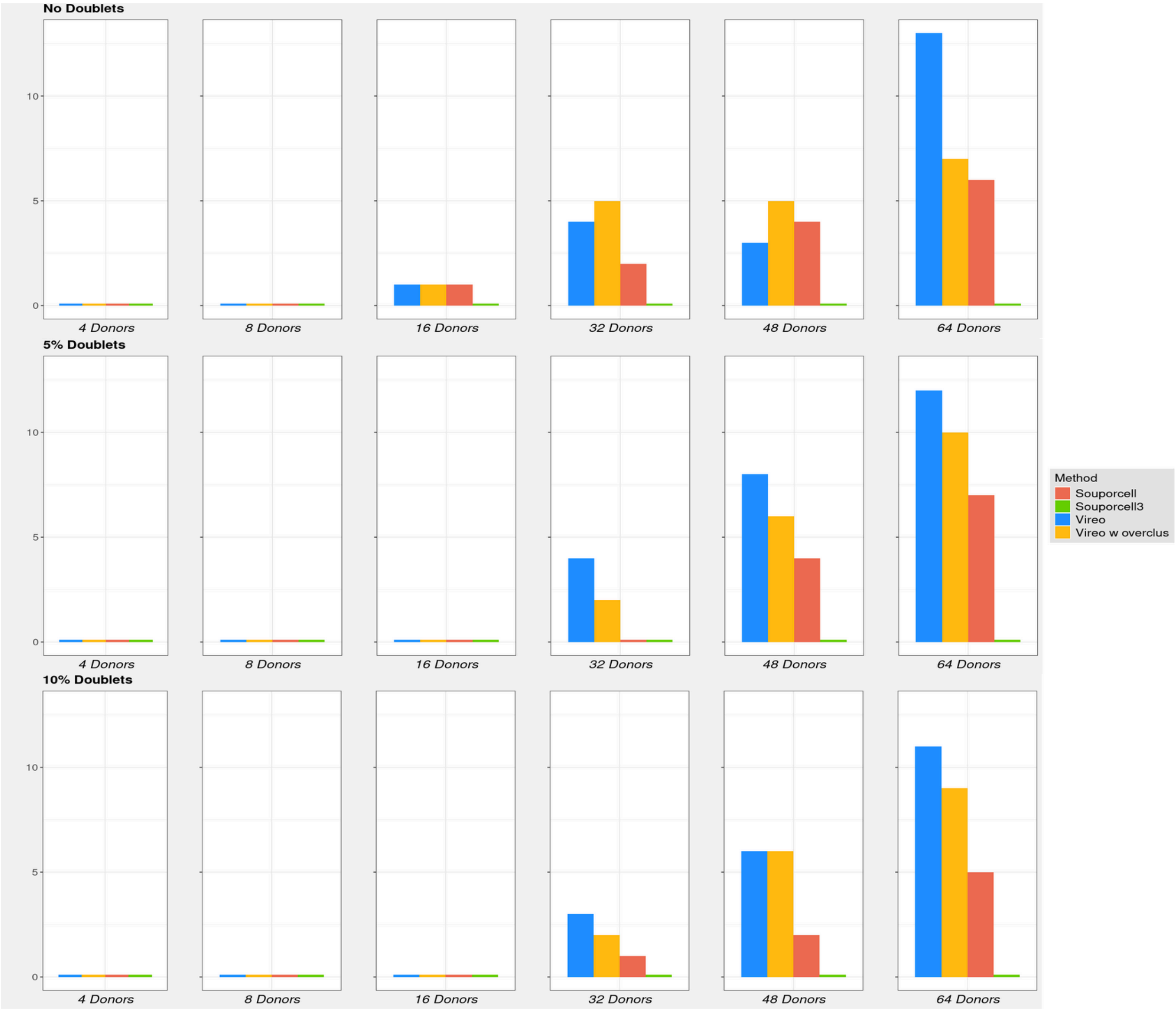


**Figure 12.** Number of duplicate donor assignments for vireo, vireo with overclustering, souporcell, and souporcell3 across different donor sizes and doublet percentages.

The Figure compares the performance of four demultiplexing tools—vireo, vireo overclus, souporcell, and souporcell3—based on the number of duplicate clusters they generate when run on datasets with varying numbers of donors (4, 8, 16, 32, 48, 64) under three conditions: no doublets, 5% doublets, and 10% doublets. Duplicate clusters indicate incorrect clustering, where cells from multiple donors are incorrectly clustered into one cluster center; thus, lower values are preferable. Under the no-doublets condition (top panel), all methods show an increasing number of duplicate clusters as the number of donors increases. vireo performs the worst, peaking at 13 duplicates at 64 donors, followed by vireo overclus and souporcell, while souporcell3 consistently yields no duplicates. With 5% doublets (middle panel), vireo performs slightly better than in the no-doublets scenario, showing 12 duplicates at 64 donors, while souporcell maintains moderate performance with only 7 duplicates. In the most challenging 10% doublet scenario (bottom panel), the pattern persists—vireo and vireo overclus exhibit a high number of duplicates, souporcell performs moderately, and souporcell3 remains the most robust, maintaining zero duplicate clusters across all donor counts. These results demonstrate that souporcell3 is the most accurate and resilient demultiplexing method across increasing dataset complexity, outperforming both vireo and earlier versions of souporcell.

#### 6.2 Adjusted Rand Index

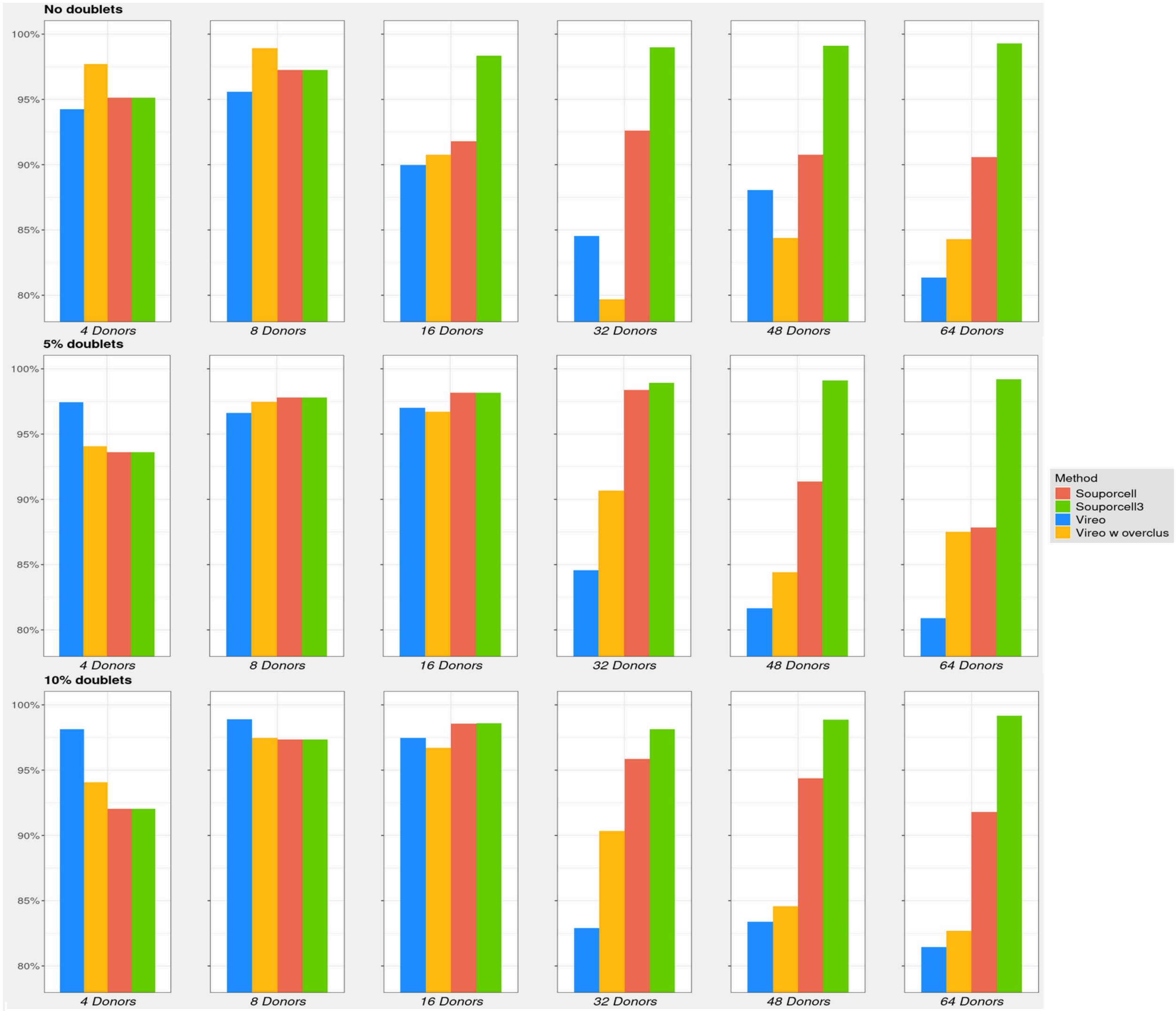


**Figure 13.** Adjusted Rand Index for vireo, vireo with overclustering, souporcell, and souporcell3 across different donor sizes and doublet percentages.

The figure compares the Adjusted Rand Index, a measure of clustering accuracy, across four demultiplexing methods—vireo, vireo overclus, souporcell, and souporcell3—on datasets with varying numbers of donors (4, 8, 16, 32, 48, 64) under three conditions: no doublets, 5% doublets, and 10% doublets. In the no-doublet condition (left panel), all methods perform well at low donor counts (4–16 donors), with Rand Index values near or above 0.95. As donor count increases to 32, 48, and 64, vireo's accuracy declines sharply, dropping below 0.86 at 48 donors and to 0.81 at 64 donors. Vireo overclus also deteriorates significantly beyond 32 donors, while Souporcell maintains better performance, and Souporcell3 consistently achieves near-perfect clustering (Rand Index > 0.99) across all donor levels. In the 5% doublet condition (middle panel), similar trends emerge: Souporcell3 remains highly accurate across all donor counts, while vireo's performance declines at 64 donors (0.82), and vireo overclus exhibits a similar drop. The 10% doublet condition (right panel) presents the greatest challenge. Souporcell3 continues to maintain a nearly perfect clustering score, while vireo experiences a pronounced decline in accuracy, especially at 64 donors (Rand Index < 0.79). Souporcell performs similarly to its results in the 5% doublet condition.

The overall trend shows that for datasets with more than 32 donors, vireo overclus outperforms vireo due to better cluster center selection strategy. However, both Souporcell and Souporcell3 still perform better in comparison. Among these, Souporcell3 consistently outshines the other tools, demonstrating the highest clustering accuracy across all dataset complexities. It maintains near-perfect Adjusted Rand Index values regardless of donor count or doublet rate, making it the most robust and reliable demultiplexing method in this comparison.

#### 6.3 Cluster maps for 10% doublet 64 donor dataset

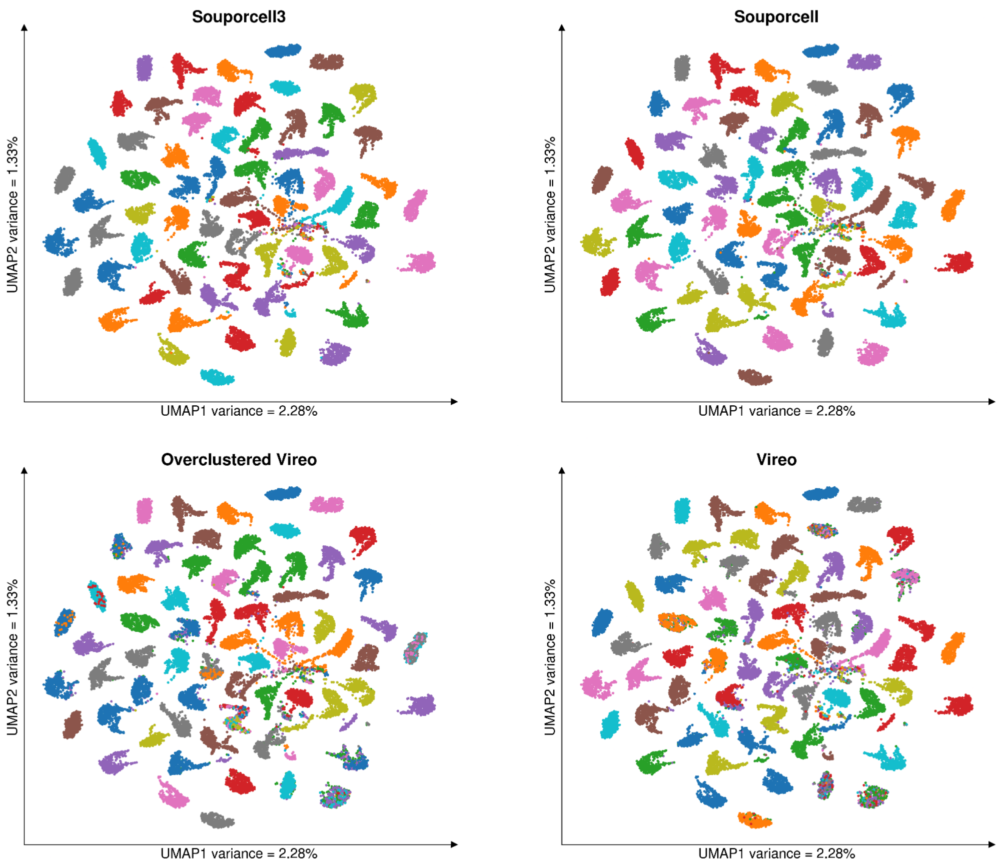


**Figure 14.** Cluster maps using UMAPs for the x and y axes, showing the result of 10% doublet, 64-donor dataset after running souporcell, souporcell3, vireo, and vireo overclus prior to doublet detection.

This UMAP plot displays the clustering of single-cell RNA-seq data from a dataset with 64 donors and 10% doublets, with doublets excluded from the visualization as it was generated before doublet detection for all methods. Each point represents a cell and is colored by its ground truth cluster (donor). The UMAP1 and UMAP2 axes capture the primary structure of the high-dimensional cluster data in two dimensions. In the souporcell3 plot, the clusters are well-separated throughout the UMAP space, indicating effective donor demultiplexing. Even in the central region, where many clusters appear close together, they remain distinct and clearly partitioned except for a few outliers, showing that the clustering algorithm is capable of resolving densely packed but separate donor profiles. In contrast, vireo and vireo overclus have many ground truth clusters where two or more clusters are assigned.

#### 6.4 Heat maps for 10% doublet 64 donor dataset

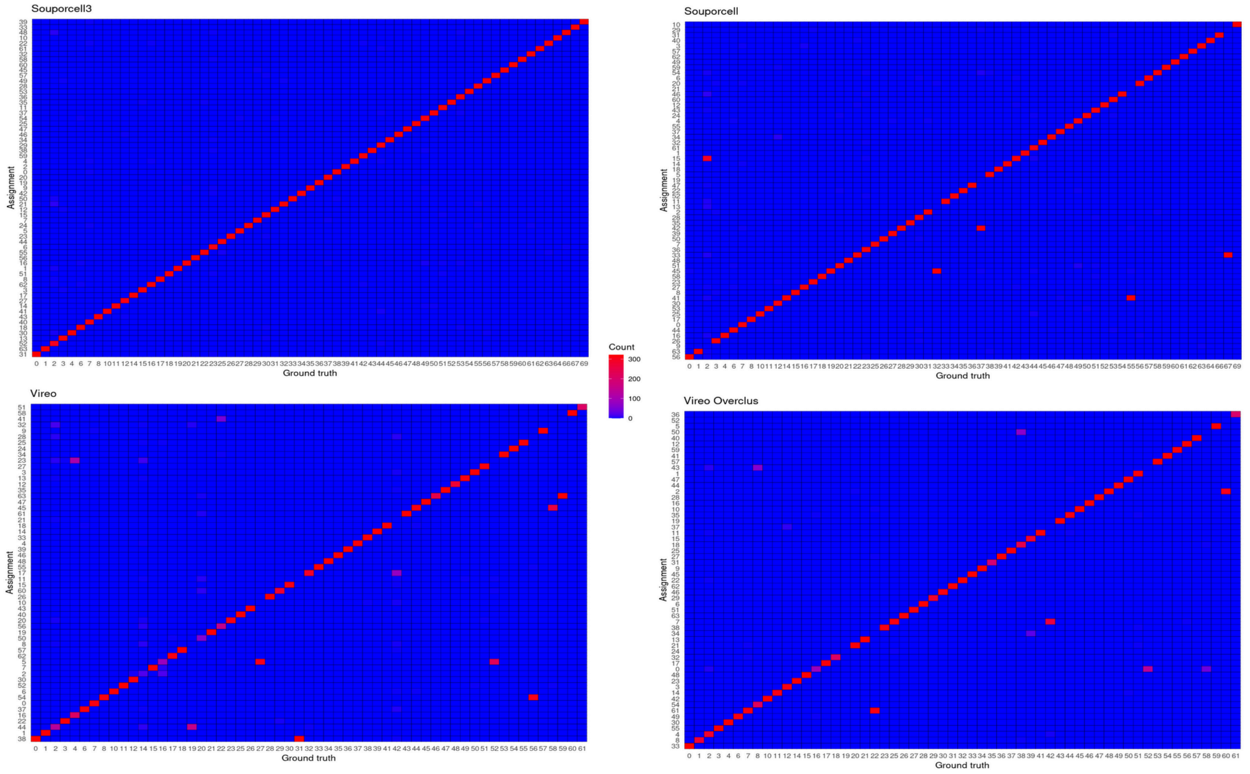


**Figure 15.** Heat maps showing cell counts for assigned cluster vs ground truth for 64 clusters of 10% doublet dataset after running with souporcell, souporcell3, vireo, and vireo overclus, prior to doublet detection.

The heat map in figure 15 displays cell counts across 64 clusters for a 64-donor dataset with 10% doublets, result generated using Souporcell3 prior to doublet detection. Each row corresponds to the cluster assignment by the method and each column represents the ground truth clusters, and the color intensity reflects the number of cells assigned, with median cell count per donor being 315. The souporcell3 heatmap shows a strong diagonal pattern, indicating that the majority of cells are correctly assigned to their respective clusters with minimal cross-contamination. This diagonal dominance demonstrates high accuracy in donor assignment, as each cluster predominantly contains cells from a single donor. The other methods all have off diagonal cell counts, meaning there are ground truth clusters where two or more clusters are assigned. The absence of significant off-diagonal cell counts further supports the effectiveness of Souporcell3 in distinguishing between donors, even in the presence of 10% doublets.
